# Assessing the multiple dimensions of poverty. Data mining approaches to the 2004-14 Health and Demographic Surveillance System in Cuatro Santos, Nicaragua

**DOI:** 10.1101/593426

**Authors:** Carina Källestål, Elmer Zelaya Blandón, Rodolfo Peña, Wilton Peréz, Mariella Contreras, Lars-Åke Persson, Oleg Sysoev, Katarina Ekholm Selling

## Abstract

We aimed to describe multiple dimensions of poverty according to the capability approach theory by applying data mining approaches to the Cuatro Santos Health and Demographic Surveillance databases, Nicaragua. Four municipalities in northern Nicaragua constitute the Cuatro Santos area, with 25,893 inhabitants in 5,966 households (2014). A local process analyzing poverty-related problems and prioritizing suggested actions, was initiated 1997 based on a community action plan 2002-2015. Priority interventions were school breakfasts, environmental protection, water and sanitation, preventive healthcare, home gardening, micro credits, technical training, stipends for university education, and the use of Internet. In 2004, a survey of basic health and demographic information was performed in the whole population followed by surveillance updates in 2007, 2009, and 2014 linking households and individuals by unique identifiers. Information included the house (floor, walls) and services (water, sanitation, electricity) as well as demographic data (birth, deaths, migration). Data on participation in interventions, on food security, household assets, and women’s self-rated health were collected in 2014. A K-means algorithm were used to cluster the household data (54 variables). The poverty ranking of household clusters using the unsatisfied basic needs index (UBN) variables changed when including variables describing basic capabilities. The households in the fairly rich cluster, having assets as motorbikes and computers were described as modern. Those in the fairly poor cluster, having different degrees of food insecurity were labeled vulnerable. Poor and poorest clusters of households were traditional e.g. in using horses for transport. Results displayed a society transforming from traditional to a modern, where the forerunners were not the richest but educated, had more working members of household, fewer children and were food secure. Those lagging were the poor, traditional and food insecure. The approach and results may be useful for an improved understanding of poverty and to direct local interventions.

## Introduction

The first of the Sustainable Development Goals aims at ending poverty in all its forms, everywhere. This is further specified as reducing by 2030 at least by half the proportion of men, women, and children of all ages that currently live in poverty <u>in all its dimensions according to national definitions</u> (our underscore) (1). This all-inclusive target addresses all dimensions of poverty, the most important determinant for health and wellbeing (2).

Poverty measures used by the World Bank and many international agencies are usually monetary measures on the national level, such as the poverty line at 1,90 purchasing power parity dollar and the Gross Domestic Product per capita. These monetary measures of poverty are possible to compare over time and across nations. In Latin America the Unsatisfied Basic Need (UBN) index has been widely used to compare poverty at the household level between different geographical areas (3, 5). UBN is a composite index that includes housing conditions, access to water and sanitation, school enrolment, education of the head of household, and the ratio of dependent household members to working age members. In the Demographic Health Surveys household asset scores have been widely used as a measurement of household socioeconomic status and poverty (5). Asset scores have been used to stratify other outcomes along a wealth axis, such as the identification and explanation of social inequalities in health (6). Asset scores cannot be used to follow or compare development over time since each index is only valid for the survey for which it was created.

The Commission on Global Poverty, assigned by the World Bank (7) has recommended the inclusion of complementary indicators when tracking poverty change over time and across settings. Further, the Commission has suggested the capability approach to poverty formulated by Amartya Sen and others as a framework to aid the development of indicators (8, 9).

The capability approach focuses on individuals and prioritizes the freedom of choice a person has over alternative lives that she or he could live (9). In this approach, the fundamental and intertwined concepts are capabilities and functions. In practice, it is often easier to evaluate achieved functions, representing the accomplished capabilities. People show adaptive preferences to their environment by adjusting their expectations to the surrounding social, cultural, political, and economic restrictions. Frequently capabilities cannot be converted to functions, thus indicating the need of equality in capabilities and functioning (10).

Amartya Sen and others have discussed whether basic capabilities should be captured in indices or decided upon by the poor themselves (8). In most cases, the basic capabilities included are adequate health, sufficient food and nutrition, adequate education to ensure basic knowledge, capability of independent thought and expression, political participation, and freedom of race, religion, and gender discrimination (10). Several indices capture multiple basic capabilities, such as the Multidimensional Poverty Index (11).

Governments have the responsibility to implement policies for poverty reduction to reach the first Sustainable Development Goal (12). Local level bottom-up interventions might, however, result in sustainable poverty reduction that might inspire decision makers at the national level. We have documented such a case in northern Nicaragua; the Cuatro Santos experiences of local poverty reduction (13). That case study showed that factors such as local ownership, locally guided multidimensional interventions, and close monitoring and evaluation of the development efforts yielded a substantial poverty reduction of household poverty from 79 to 47 % over a ten year period (2004-14) (13).

In the Cuatro Santos area, a Health and Demographic Surveillance System (HDSS) was established in 2004 with the latest update in 2014. Participation in micro credit programs, the involvement of young individuals in technical training, and home gardening were associated with the transition of households out of poverty (14). The Unsatisfied Basic Needs scoring of households was used to identify geographic areas with higher levels of poverty to target interventions (13). However, poverty indices, such as the Unsatisfied Basic Needs or asset scores, have limitations for a context-specific description of poverty. To address this problem, a data mining method, a variant of the K-means clustering algorithm (15), is an alternative approach to identify patterns, which might describe poverty in a local context in a multidimensional way. Thus, this paper’s aims to describe the multiple dimensions of poverty according to the capability approach theory by using data mining approaches to the Cuatro Santos Health and Demographic Surveillance databases, Nicaragua.

## Methods

### Study setting, population, and design

The Cuatro Santos area, situated in the northern part of Chinandega, Nicaragua, consists of four municipalities of similar population size. In 2014 totally 25,893 inhabitants lived in 5,966 households in an area located 250 km northwest of the capital of Nicaragua, Managua, in a mountainous terrain bordering Honduras. The climate is predominantly dry and the traditional sources of income have been the cultivation of grains and raising livestock, now with an increasing number of small-scale enterprises. This area was strongly affected by the Contras war in the 1980s and the hurricane Mitch in October 1998. Since that time a significant proportion of the population has out-migrated due to economic reasons, including fixed or seasonal work or search for employment (16).

### Community interventions in Cuatro Santos

Starting in 1997, representatives of the four municipalities, the local non-governmental organizations, local government leaders, and representatives of national institutions initiated a process labeled “decoding reality”, which was inspired by Paulo Freire (17). This process included an analysis of the local poverty-related problems, prioritization among suggested actions, and an action plan that was approved as the Cuatro Santos Area Development Strategy 2002 to 2015. This strategy aimed at efforts to develop the area by use of local resources, informed by data from the surveillance system, and to attract international cooperation. The concepts of local ownership and participation were central, and the efforts included consensus decision-making and reconciliation in case of conflicts. Priority interventions were school breakfasts, environmental protection, water and sanitation, preventive healthcare, home gardening, micro credits, technical training, stipends for university education, and telecommunications including access to and training in the use of Internet. Data collection through a Health and Demographic Surveillance System was central for monitoring of trends over time, and research evaluation of various aspects (13,14).

### Cuatro Santos Health and Demographic Surveillance System

In 2004, a census and cross-sectional data collection of basic health and demographic information was performed in the whole population. Follow-up surveys were performed in 2007, 2009, and 2014. Unique identifiers of households and individuals linked the data. Demographic changes in the households, such as birth, death, and migration, were registered. Household data included information on the house (floor, walls) and services (water, sanitation, electricity). All women aged 15– 49 years living in the households provided retrospective reproductive histories (14). In the 2009 and 2014 updates, questions were included on participation in the following interventions: access to water and latrines, micro credit, home gardening, technical education, school breakfast programs, and telecommunications. In the 2014 update, data on food security, household assets, and women’s self-rated health were collected. For the present study, data from the 2014 update including data on earlier events and interventions were used.

Fieldwork conducted by local female fieldworkers was carefully supervised, printed forms were checked before computerization, and the forms were returned to the field if the information was missing or suspected to be incorrect. Further data quality controls were completed after computerization including logical controls. Data were carefully cleaned and stored in a relational database (Microsoft Access 2007®).

### Variables (see Table 1)

Persons residing in a household at the time of the field survey defined the household. Migration was defined as a household member aged 18–65 who migrated in or out of the household since the previous update (5 yrs.). The Unsatisfied Basic Needs index (16) was composed by four components: (I) housing conditions (unsatisfied: walls of wood, cardboard, plastic or earthen floor); (II) access to water and latrine (unsatisfied: water from river, well, or bought in barrels and no latrine or toilet); (III) school enrolment of children (unsatisfied: any children 7–14 years of age not attending school); and (IV) education of head of the family and ratio of dependent (<15 yrs. and >65 yrs.) household members working-age members (15-65 yrs.) (unsatisfied: head of the family illiterate or dropped out of primary school and ratio of dependent household members working-age members > 2.0). Each component rendered a score of zero, if satisfied, and one, if unsatisfied. Thus, the total sum varied from zero to four. Households with zero or one unsatisfied basic needs were considered non-poor, while poor households had two to four unsatisfied basic needs. Characteristics of houses and households were also included in the cluster analyses, such as the material of walls, floor, access to electricity, type of stove, access to water, and type of toilet. The interventions implemented in the area were represented by household-related information on such participation. The presence of a water meter indicated that the household had got water installed as part of the last decade’s interventions. Also, information was included on previous and current participation in home gardening, if anyone in the household had received micro credits or had participated in technical training.

**Table 1.**
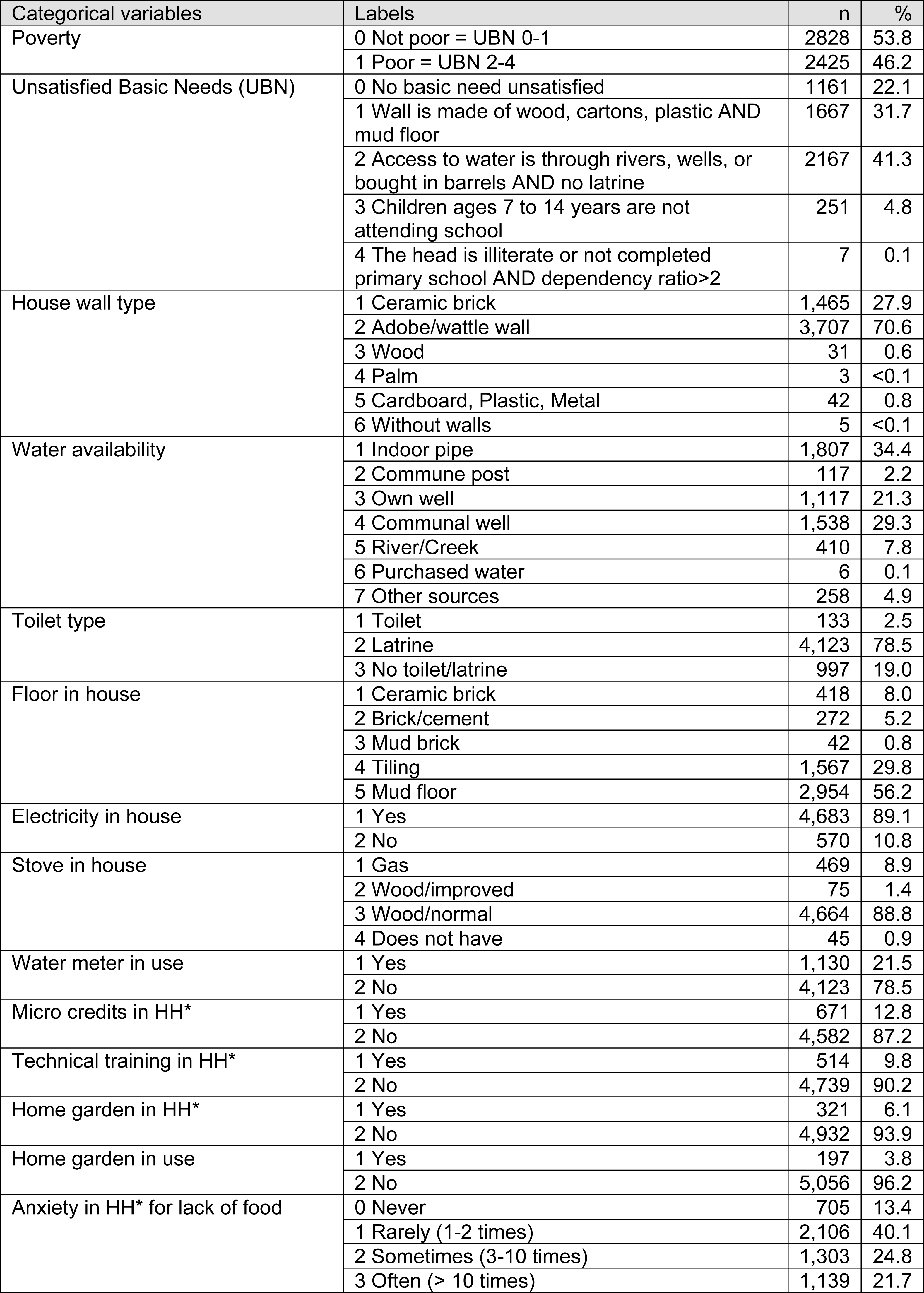
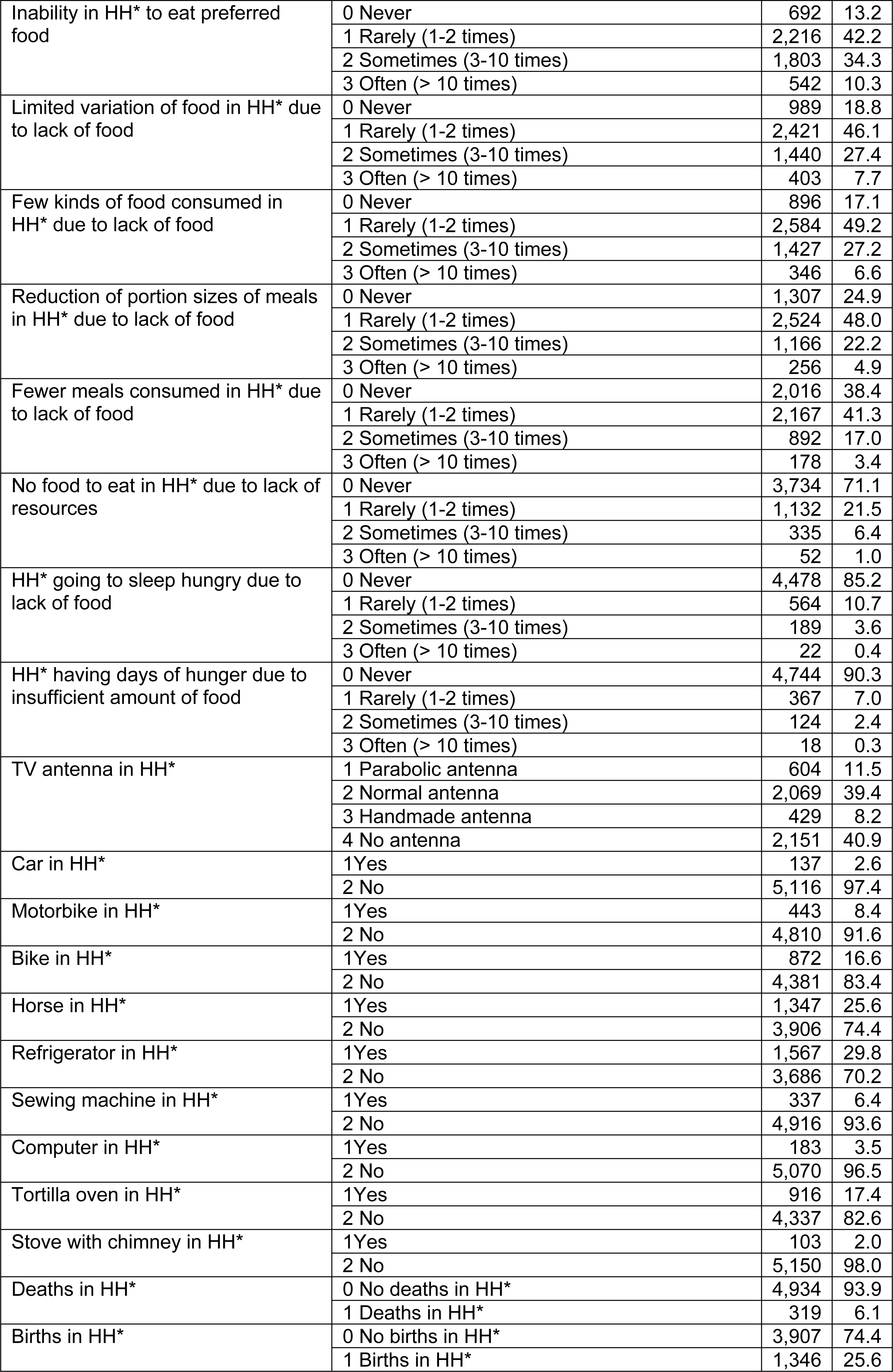
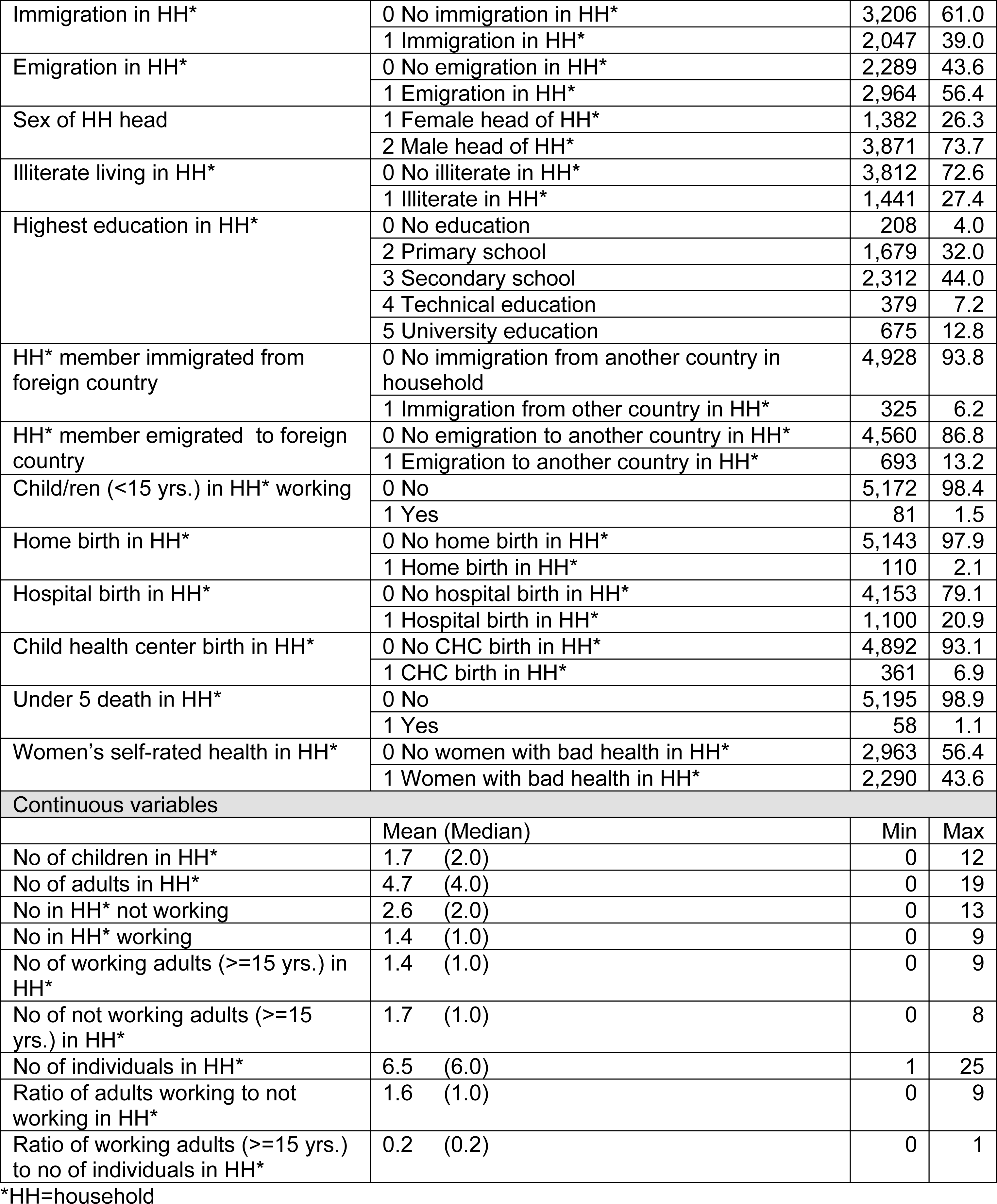
List of variables included in the analyses of Cuatro Santos database, Nicaragua 2014, including descriptive statistics.

A nine-item Household Food Insecurity Access Scale (HFIAS), version 3, was used (18). The respondents were either the head of the household or the person responsible for the household expenditure and food preparation during the last four previous weeks. This scale covers experiences regarding 1) anxiety in the household due to lack of food; 2) inability to eat preferred food because of lack of resources; 3) limited variety of food due to lack of resources; 4) consumption of few kinds of food because of lack of resources; 5) reduction of portion sizes of meals due to lack of food; 6) consumption of fewer meals per day because of lack of food; 7) no food to eat in the household because lack of resources; 8) going to sleep at night hungry due to lack of food, and 9) days of hunger because of insufficient amounts of food to eat. For each affirmative answer, the person provided additional information on the frequency in a four-point scale (never, rarely, sometimes, often).

Household assets were TV antenna, car, motorbike, bike, horse, refrigerator, sewing machine, computer, tortilla oven, and a chimney for the wood-burning stove.

The individual variables collected 2014 were derived and aggregated at the household level, and thereafter merged with the variables at household level. We constructed variables on births and deaths in the household during the recent update period, also including information on under-5 death, number of adults and children living in the household, number of adults and children working, number of adults not working, and the ratio between adults working and not working, as well as the ratio between adults working and number of individuals in the household. Further, data were included on in-and out-migration, including from foreign countries, gender of household head, any illiteracy, and the highest education level in the household (none, primary, secondary, technical, university education). Information was also included if a home-, health center-, or hospital birth had happened since the last update (5yrs).

Women’s self-rated health was assessed for all resident women of reproductive age (15–49 years) at time of the interview by a five-point Likert scale based on the following question: “In general, how would you assess your health today?” The interviewer provided the following options: very good, good, medium, bad, or very bad. This information was classified as good (very good, good, medium) or bad (bad, very bad) health. No household had a mix of good and bad self-assessed health when aggregating this information to household level. The total dataset included 54 variables.

### Analytical methods

All analyses were performed on the household level. The variables included are displayed in Table 1. A variant of the K-means clustering algorithm (15) called SimpleKMeans in Weka (19) was used to perform a clustering of our data. The reason for choosing K-means algorithm was that K-means is “the most popular and the simplest partitional algorithm” (20). The K-means algorithm computes K points called centroids and then assigns the data points to their respective closest centroid. This leads to forming K groups (clusters) of observations in the data where observations within each cluster have similar properties. To evaluate the quality of the clustering, data were split into training and test sets. Cluster centroids were computed from the training data and tested on the test data by using the closest-centroid-principle. Properties of the training and test clusters were compared and the robustness was evaluated.

Categorical variables were transformed into dummy variables and included in the K-means cluster analysis and after being scaled, the numerical variables were also included in the analysis. Repeated analyses where performed forcing data into 2 to 10 clusters. Default values were taken for all other settings of the algorithm. A so-called scree plot was created displaying cluster Sums of Squared Errors (y-axis) and number of clusters (x-axis) (S2 Figure, Supplemental Figure 1). An appropriate number of clusters in the plot can be found by identifying the level of the x variable where the saturating starts. Six clusters were selected after inspection of this scree plot and checking cluster sizes. The Euclidian distance was applied and the data were randomly split into training (66 %) and test (44 %) sets. The meaning of the clusters was interpreted by evaluating the cluster centroids (percentages for dummy variables of categorical variables and averages for numerical variables) in each cluster in relation to each other and to the full data.

**Fig 1.**
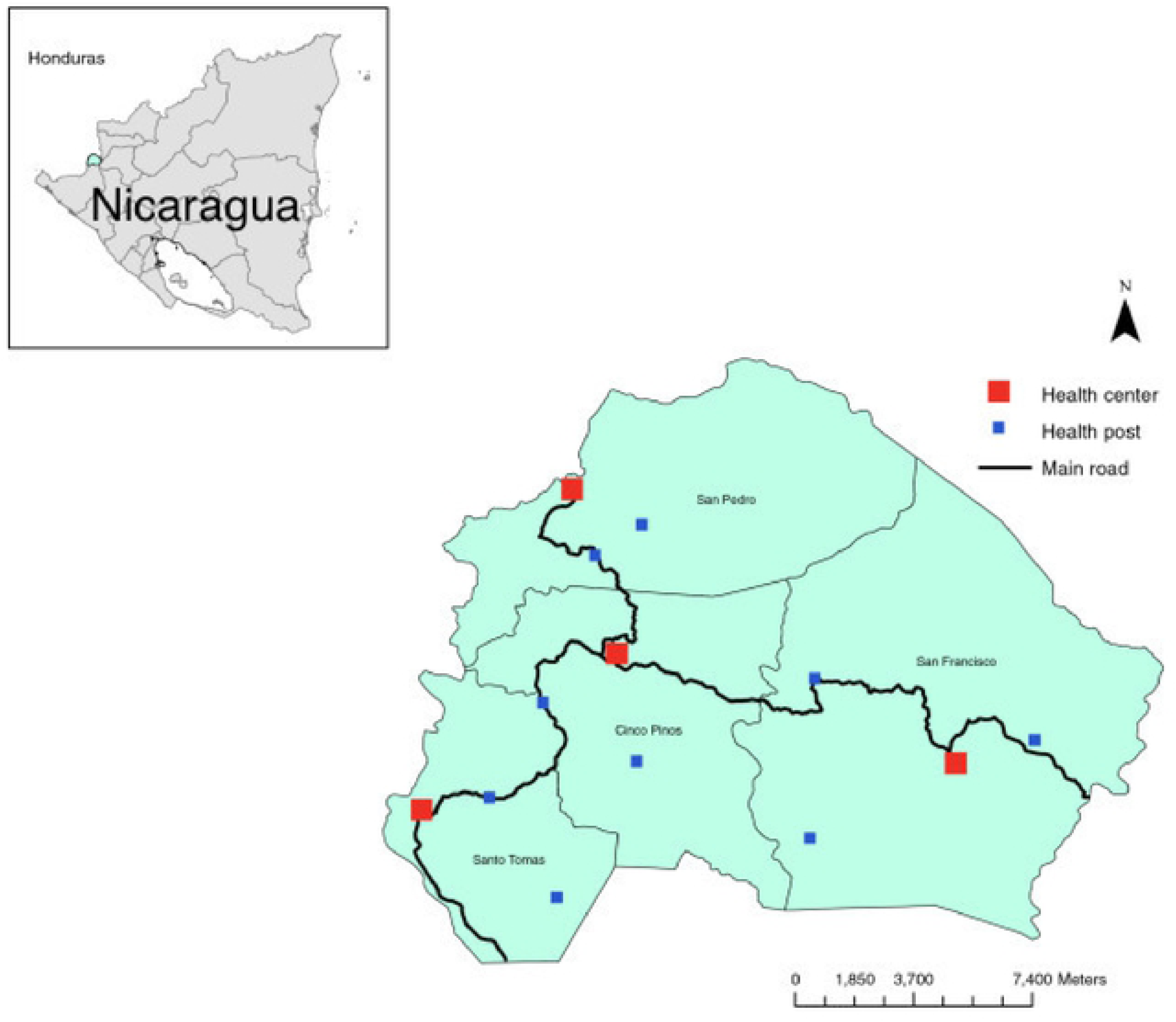
The Cuatro Santos area showing the four municipalities and health facilities. The area is marked in the inserted Nicaragua map.

Variable groups of categories were analyzed in a stepwise order to generate an assessment of poverty. These categories were included in the following order: a) poverty assessed by the variables poverty and UBN and variables in UBN except head of household’s education, children’s school enrolment, and ratio dependents to working household members, b) assets, c) food insecurity, d) interventions, e) derived individual variables (see Table 1 for included variables, and Supplemental Table 1 for full cluster analysis output where the categories are color marked). The emerging patterns were evaluated and the clusters were labeled in words as reported in results. Table 2 shows the essential variables extracted from Supplemental Table 1, yielding the labeling words.

**Table 2.**
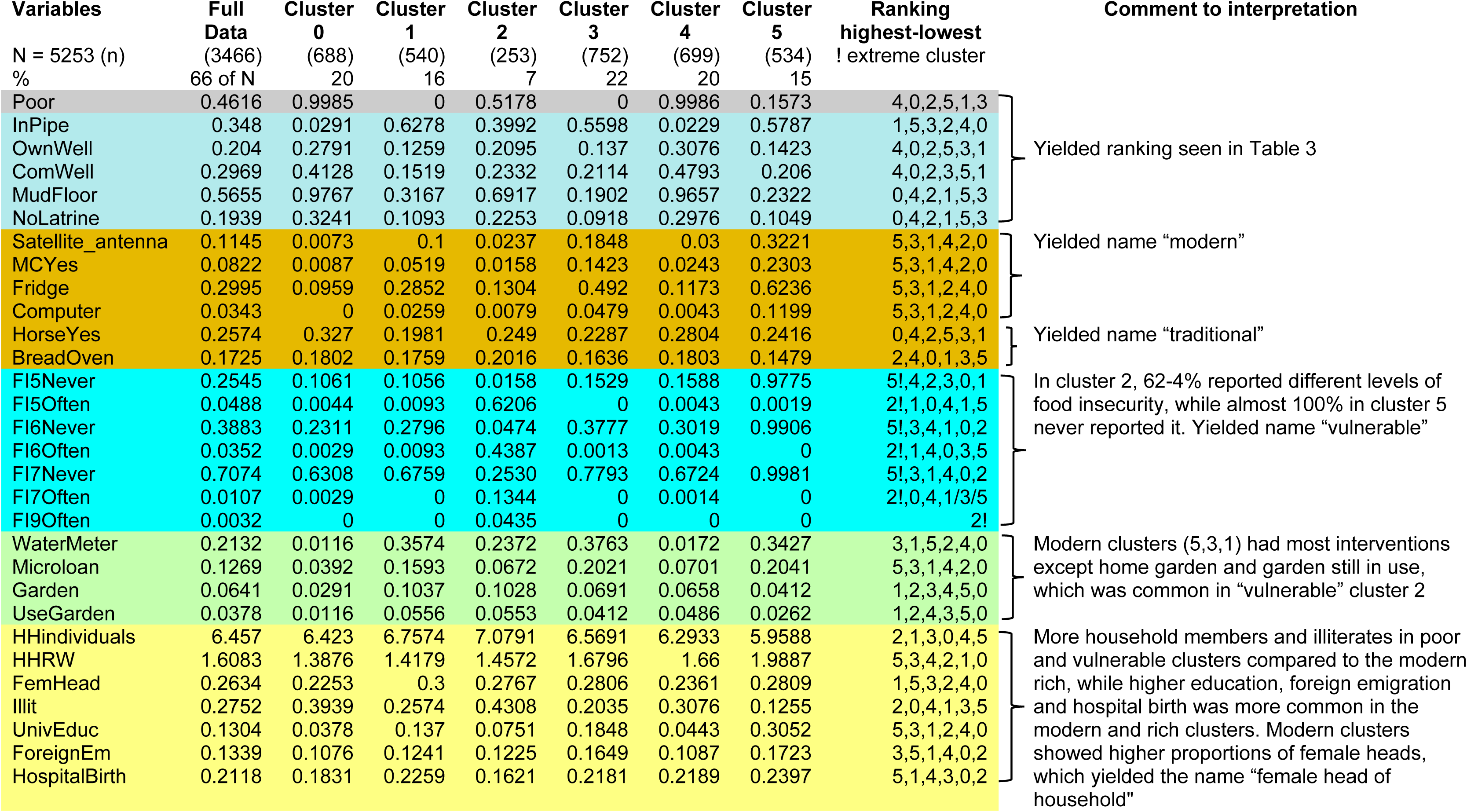
Meaningful variables used in the analysis of clusters illustrating naming of clusters. (Extracted from S1 Appendix, Supplemental Table 1, categories color marked as follows: Grey= Poverty assessed by the variable poverty, Light blue = variables in UBN, except head of household’s education, children’s school enrolment, and dependency ratio, Dark yellow = assets, Turquoise = food insecurity, Green = interventions, Light Yellow = derived individual variables)

### Ethical considerations

The information was collected as part of the Health and Demographic Surveillance update survey in 2014. The Ethical Review Board of Biomedical Research at the National Autonomous University of León approved the HDSS data collection (FWA00004523/IRB0000334 ACTA No. 81). Informed verbal consent was obtained from the participants. They were free to end their participation at any time. Data were stored in a safe electronic platform with an alphanumeric identification number instead of names of participants to protect confidentiality.

## Results

Of the 5,966 households included in the 2014 update of the HDSS, 5,253 (88 %) were included in the analyses after eliminating households with missing values on any variable. The major reasons to omissions were houses included in the database as households while, in fact, not being living quarters, e.g., schools, health centers, or abandoned houses. Included data measured experiences since the last update (5 yrs.) and earlier participation in interventions. The basic characteristics of the households are shown in Table 1.

### Cluster analyses

The patterns emerging from the variables separating the clusters the most (in the following text these variables are called essential variables), extracted from S1 Appendix, Supplemental Table 1, and the labeling of clusters is illustrated in Table 2.

Poverty assessed by the first category, i.e., the dichotomized variable poverty and the 5 UBN categories (0-4) and the variables characterizing the household physical conditions and the water and sanitation conditions (S1 Appendix, Supplemental Table 1, and Table 2) yielded a ranking by <u>poverty status</u> as shown in Table 3 with essential variables being *poor, water source, mud floor* and *no latrine.*

**Table 3.** Results from cluster analysis of first ranking using Unsatisfied Basic Needs (UBN) variables from the Health and Demographic Surveillance System, Cuatro Santos, Nicaragua.

| Cluster<br>(% of HH<sup>2</sup>) | Poverty<sup>1</sup> |
| --- | --- |
| 4 (20%) | Poorest |
| 0 (20%) | Poor |
| 2 (7%) | Fairly poor |
| 5 (15%) | Fairly rich |
| 1(16%) | Rich |
| 3 (22%) | Richest |

1. Rich and poor refers to our UBN categories and household characteristics included in the UBN 2. HH=households

Cluster 5 (Table 3) showed to be the most <u>modern</u> cluster having assets that were modern equipment like satellite dish antenna, computer, refrigerator, motorbike. Clusters 3 and 1 had also these assets but to a lesser extent. Clusters 0, 2, and 4 were more <u>traditional</u> with assets as horses and tortilla bread ovens in higher proportions, illustrating that transportation and earnings of living by selling tortillas were carried out as in earlier times. These assets yielded the names <u>traditional</u> and <u>modern</u>.

The distribution of food insecurity variables showed that cluster 2 (7% of households) was far more food insecure than all other clusters including all aspects of food security and that cluster 5 was food secure. These characteristics added the descriptive word <u>vulnerable</u>.

The most modern, richest and least vulnerable cluster had participated most in interventions. One exception was home gardening and still using a garden, which was more common among the traditional, and vulnerable clusters, especially the food insecure cluster 2. The latter intervention had however, reached few households. The essential variables were water meter, micro credit, technical training and home gardening.

When including all variables, the re-ranking displayed clusters of <u>multidimensional poverty</u> and the derived individual variables made this new ranking more distinct (Table 4). More household members and children were found in poor and vulnerable clusters compared to the modern rich, while higher education was more common in the modern and rich clusters. Overall, female and male-headed household proportions were ¼ and >, respectively and the more modern clusters showed higher proportions of female heads, which rendered the descriptive word <u>female head of household</u> in naming of clusters. The following were the most essential of the derived individual variables; number of household individuals, ratio of adults working to those not working, female/male household head, illiterate individuals in household, university education in household, foreign emigration in household, and hospital birth, which all strengthened the multidimensional poverty group ranking and modern or traditional labeling.

**Table 4.** Results from cluster analysis second ranking including all variables from the Health and Demographic Surveillance System, Cuatro Santos, Nicaragua.

| Cluster<br>(% of HH<sup>2</sup>) | Multidimensional poverty<sup>1</sup> |
| --- | --- |
| 2 (7%) | Fairly poor, most vulnerable, fairly traditional |
| 0 (20%) | Poor, traditional |
| 4 (20%) | Poorest, traditional |
| 1 (16%) | Rich, fairly modern, female head of household |
| 3 (22%) | Richest, fairly modern, female head of household |
| 5 (15%) | Fairly rich, most modern, female head of household |

1. Rich and poor refers to our Unsatisfied Basic Needs (UBN) categories and household characteristics included in the UBN, while modern and traditional refer to assets, interventions, number of adults and children in household, education, emigration, and hospital births. Vulnerable refers to food security and female head of household to proportion of female-headed households 2. HH=households

## Discussion

This study is unique as it assesses multidimensional poverty using data at household level with a large number of variables taking advantage of a data mining technique. Variables assessing household conditions, food insecurity, access to interventions, demographic and mortality events were used. We found six clusters of households with differences between them, and with similarities within them, based on their shared variables.

The ranking of households using the unsatisfied basic needs index (UBN) variables in the cluster analysis were changed when including more variables describing basic capabilities. Most importantly, the fairly rich cluster 5 showed to be the most modern, with modern assets such as motorbikes and computers. The fairly poor cluster 2 showed to be the most vulnerable, having varying degrees of food insecurity, something that the most modern cluster never experienced. The poor and poorest clusters were traditional, illustrated by the use of horses for transport. Men headed two-thirds of households, but the proportion headed by women were higher among the modern rich. Altogether, the results pointed at a traditional society in transition to becoming modern. The forerunners were educated, had more working members in the household, had fewer children and were food secure but were not the richest according to the Unsatisfied Basic Needs characteristics. While those lagging were the poor, traditional and food insecure.

The importance of food insecurity was illustrated by the fairly poor becoming the most vulnerable in the multidimensional poverty analysis. It should be noted that the finding that participation in interventions, as for instance getting water installed, receiving a microloan, or engaging in technical training coincides with better welfare.

The Health and Demographic Surveillance data have been judged to be of high quality (13, 14) and covered the whole population in the Cuatro Santos area with very few non-participants, thereby providing a reliable basis for analyzes. The temporality of poverty predictors (a predictor happening before poverty) was not fully captured by our design. Based on the dates of the initiation of the interventions stored in our database, however, we can state that most interventions happened before the 2014 survey. The timing of acquisition of assets was neither known, nor did we know when the head of household was established, although analyses have shown stability over time of household head. Food insecurity answers covered experiences during the last four weeks before survey.

Cluster analysis is a powerful method to identify hidden groups in the data, and K-means is an algorithm, which is fast, simple to use and interpret. Compared to some other clustering methods, number of clusters can be visually selected on the scree plot. It is worth mentioning that the Euclidian distance was used, in which categorical variables were transformed to dummy variables and the continuous variables were scaled. These metrics are very general and do not rely on any application assumptions. Our cluster analysis has, however, some limitations. Firstly, K-means clustering optimizes the distances to the cluster centroids which means that spherical clusters are relatively easy to detect but if a cluster has a complicated shape, K-means clustering might split this into two or more parts. Secondly, all variables were included in the distance measure of the cluster analysis, including potentially irrelevant variables. This might in theory lead to blurring of some clusters, although in our analysis, we managed to obtain well-interpretable clusters with clearly distinct properties.

The interpretation and the choice of descriptive names of clusters was a subjective exercise that depend on the analyst’s pre-understanding. The naming can, however easily be reviewed by studying Supplemental Table 2 which displays the cluster analysis.

Food insecurity is essential for wellbeing as shown in the multidimensional analysis of poverty. This was also reflected in the association between low self-rated health and food insecurity in a previous study from our group using data from the same surveillance system (21).

Interventions, such as water installation, micro credits, and participation in educational activities, positively influenced welfare, confirming our earlier results (14).

The randomized controlled trials evaluation of multifaceted programs in six countries have comparison villages (22) and a recent publication tried to accomplish comparisons for the Millennium development villages evaluation (23), both reporting positive results for complex interventions aiming for increased welfare in poor areas. The Cuatro Santos case study (13) has no comparison area so we cannot rule out that the general transformation of the Nicaraguan society is a reason for the improvements in welfare seen in the area. The finding in this analysis of multiple dimensions of poverty do however, provide some support that the interventions contributed to poverty reduction.

The Health and Demographic Surveillance data did not cover all aspects of basic capabilities. Even so, we consider having captured the multidimensionality of poverty stressed by the capability approach. We would like to argue that the results were meaningful, comprehensible and familiar in the area, based on a feedback and inference discussion held in the area with local community leaders and representatives of different sectors of society including health and security as well as lay people from the communities. These local community representatives confirmed the usefulness of this and similar further analyses for targeting interventions intending to reduce inequity.

## Conclusion

The classification of households from rich to poor based on the unsatisfied basic needs assessment was modified by a multidimensional analysis of poverty. The “fairly rich” households based on the unsatisfied basic needs index were the forerunners of modern lifestyle with higher welfare, while the fairly poor were the most food insecure. Results obtained from a cluster analysis may be useful for increased understanding of poverty. Health and Demographic Surveillance data, maybe enhanced by computer applications, could be analyzed and guide priority setting and direct interventions to increase general welfare.

## Supporting information

### S1 Appendix

Supplemental Table 1. Cluster analysis output with the categories color marked as follows: Grey= Poverty assessed by the variables poverty and Unsatisfied Basic Needs (UBN), Light blue = variables in UBN, except head of household’s education, children’s school enrolment, and dependency ratio, Dark yellow = assets, Turquoise = food insecurity, Green = interventions, Light Yellow = derived individual variables.

### S2 Figure

Supplemental figure 1. Scree plot displaying within cluster Sums of Squared Errors (y-axis) and number of clusters (x-axis) from K-means cluster analysis of data from Cuatro Santos Health and Demographic Surveillance System, 2014

## References

1. UN. The Sustainable Development Goal 1. Accessed 5.29.18 from: https://sustainabledevelopment.un.orgsdg

2. OECD. Economic well-being. In: OECD Framework for Statistics on the Distribution of Household Income, Consumption and Wealth. 2013. pp. 1–15.

3. Hammill M. Income poverty and unsatisfied basic need. Mexico City: ECLAC; 2009 Dec 10.

4. Peña, R., Pérez, W., Meléndez, M., Källestål, C., Persson, L.-Å. The Nicaraguan Health and Demographic Surveillance Site, HDSS-Leon: a platform for public health research. Scand J Public Health 2008;36: 318–25. doi:10.1177/1403494807085357

5. Howe LD, Galobardes B, Matijasevich A, Gordon D, Johnston D, Onwujekwe O, et al. Measuring socio-economic position for epidemiological studies in low- and middle-income countries: a methods of measurement in epidemiology paper. International Journal of Epidemiology. 2012 Jul 13;41(3):871–86. doi:10.1093/ije/dys037

6. Barros AJ, Ronsmans C, Axelson H, Loaiza E, Bertoldi AD, França GV, et al. Equity in maternal, newborn, and child health interventions in Countdown to 2015: a retrospective review of survey data from 54 countries. The Lancet. 2012 Mar 31;379(9822): 1225–33. doi:10.1016/S0140-6736(12)60113-5

7. World Bank. Monitoring Global Poverty: Report of the Commission on Global Poverty. Washington, DC: World Bank; 2016 Nov pp. 1–263. doi:10.1596/978-1-4648-0961-3.

8. Clark DA. The Capability Approach: Its Development, Critiques and Recent Advances GPRG-WPS-032. 2005; Dec 21: 1–18.

9. Sen A. “Justice: Means versus Freedoms…” Philosophy Public Affairs. 1990;19(2): 111–21.

10. Cosgrove S, Curtis B. Understanding global poverty. London and New York: Routledge; 2018.

11. Alkire S, Santos ME. A Multidimensional Approach: Poverty Measurement & Beyond. Soc Indic Res. 2013 Feb 13;112(2): 239–57.

12. Días Langou G, Florito J. Starting strong. Implementation of social SDGs in Latin America. Overseas Development Institute, Southern Voice on Post-MDG International Development Goals, Gala Diaz Langou, Florito J, editors. 2016 Dec pp. 1–30.

13. Blandón EZ, Källestål C, Peña R, Pérez W, Berglund S, Contreras M, et al. Breaking the cycles of poverty: Strategies, achievements, and lessons learned in Los Cuatro Santos, Nicaragua, 1990–2014. Global Health Action. 2017;10: 1–12.

14. Pérez W, Zelaya Blandón E, Persson L-Å, Peña R, Källestål C. Progress towards millennium development goal 1 in northern rural Nicaragua: Findings from a health and demographic surveillance site. Int J Equity Health. 2012;11: 43.

15. Lloyd S. Least squares quantization in PCM. IEEE transactions on information theory. 1982 Mar;28(2): 129–37.

16. Gustafsson C. For a better life. PhD Thesis, Umeå University. 2014. pp. 1–358. Available from: http://umu.diva-portal.org/

17. Au W. The dialectical materialism of Paulo Freire’s critical pedagogy. REA. 2017 Aug 23;25(2): 171–95.

18. Ballard TJ, Kepple AW, Cafiero C. The Food Insecurity Experience Scale. 2013. Technical Paper. Rome, FAO. (available at http://www.fao.org/economic/ess/ess-fs/voices/en/).

19. Witten IH, Frank E, Hall MA, Pal CJ. The WEKA Workbench. 4 ed. San Francisco: Morgan Kaufmann Publishers Inc; 2016.

20. Jain, A. K. Data clustering: 50 years beyond K-means. Pattern recognition letters. 2010; 31(8): 651–66.

21. Pérez, W., Contreras M, Peña, R, Zelaya Blandón, E., Persson, L.-Å., Källestål, C. Food insecurity and self-rated health in rural Nicaraguan women of reproductive age: a cross-sectional study. Journal for Equity in Health 2018;17: 146. doi.org/10.1186/s12939-018-0854-5

22. Banerjee A, Duflo E, Goldberg N, Karlan D, Osei R, Pariente W, et al. A multifaceted program causes lasting progress for the very poor: Evidence from six countries. Science. 2015 May 14;348: 1260799–9.

23. Mitchell S, Gellman A, Ross R, Chen J, Bari S, Huynh UK, et al. The Millennium Villages Project: a retrospective, observational, endline evaluation. The Lancet Global Health. 2018 Mar 30;6: e500–13.

